# Comparison of the human gastric microbiota in hypochlorhydric states arising as a result of *Helicobacter pylori*-induced atrophic gastritis, autoimmune atrophic gastritis and proton pump inhibitor use

**DOI:** 10.1101/144907

**Authors:** Bryony N. Parsons, Umer Zeeshan Ijaz, Rosalinda D’Amore, Michael D. Burkitt, Richard Eccles, Luca Lenzi, Carrie A. Duckworth, Andrew R. Moore, Laszlo Tiszlavicz, Andrea Varro, Neil Hall, D. Mark Pritchard

## Abstract

**Objective:** Several conditions associated with reduced gastric acid secretion confer an altered risk of developing a gastric malignancy. *Helicobacter pylori*-induced atrophic gastritis predisposes to gastric adenocarcinoma, autoimmune atrophic gastritis is a precursor of type I gastric neuroendocrine tumours, whereas proton pump inhibitor (PPI) use does not affect stomach cancer risk. We hypothesised that each of these conditions was associated with specific alterations in the gastric microbiota and that this influenced subsequent tumour risk.

**Design:** 95 patients (in groups representing normal stomach, PPI treated, *H. pylori* gastritis, *H. pylori*-induced atrophic gastritis and autoimmune atrophic gastritis) were selected from a cohort of 1400. RNA extracted from gastric corpus biopsies was analysed using 16S rRNA sequencing (MiSeq).

**Results:** Samples from normal stomachs and patients treated with PPIs demonstrated similarly high microbial diversity. Patients with autoimmune atrophic gastritis also exhibited relatively high microbial diversity, but with samples dominated by *Streptococcus. H. pylori* colonisation was associated with decreased microbial diversity and reduced complexity of co-occurrence networks. *H. pylori*-induced atrophic gastritis resulted in lower bacterial abundances and diversity, whereas autoimmune atrophic gastritis resulted in greater bacterial abundance and equally high diversity compared to normal stomachs. Pathway analysis suggested that glucose-6-phospahte1-dehydrogenase and D-lactate dehydrogenase were over represented in *H. pylori*-induced atrophic gastritis versus autoimmune atrophic gastritis, and that both these groups showed increases in fumarate reductase.

**Conclusion:** Autoimmune and *H. pylori*-induced atrophic gastritis were associated with different gastric microbial profiles. PPI treated patients showed relatively few alterations in the gastric microbiota compared to healthy subjects.

**SIGNIFICANCE OF THIS STUDY:** *What is already known about this subject?:*

- Some conditions which result in reduced gastric acid secretion and hypochlorhydria are associated with an increased risk of gastric tumourigenesis.
- This risk is different in patients with *H. pylori*-induced atrophic gastritis, autoimmune atrophic gastritis and chronic proton pump inhibitor use.
- Hypochlorhydria and *H. pylori* infection cause alterations in the composition of the gastric microbiota.

*What are the new findings?:*

- We used 16S rRNA sequencing to characterise the microbiota in gastric corpus biopsies from a well characterised cohort of patients.
- The gastric microbiota was different in patients who were hypochlorhydric as a result of *H. pylori*-induced atrophic gastritis, autoimmune atrophic gastritis and proton pump inhibitor use.
- Biochemical pathways associated with gastric carcinogenesis such as the fumarate reductase pathway were predicted to be altered in patients with atrophic gastritis.

*How might it impact on clinical practice in the foreseeable future?:*

- Understanding how the microbiota that colonise the hypochlorhydric stomach influence gastric carcinogenesis may ultimately permit stratification of patients’ subsequent tumour risk.
- Interventions that alter the composition of the gastric microbiome in hypochlorhydric patients with atrophic gastritis should be tested to investigate whether they alter the subsequent risk of developing gastric malignancy.

## INTRODUCTION

Gastric adenocarcinoma is the third most common cause of cancer related mortality worldwide[1] and most cases are associated with chronic *Helicobacter pylori* infection. Gastric cancer usually develops via the premalignant condition of gastric atrophy, which is associated with the loss of acid-secreting parietal cells[2]. The resulting hypochlorhydria potentially leads to alterations in the composition of the gastric microbiota by providing a more favourable environment for colonisation. It is currently unclear to what extent the non-*H. pylori* gastric microbiota contributes towards gastric carcinogenesis. Although the hypochlorhydria associated with autoimmune atrophic gastritis also increases the risk of developing gastric adenocarcinoma[3], it is more frequently associated with the development of another tumour, the type I gastric neuroendocrine tumour (NET)[4]. However, hypochlorhydria does not always increase the risk of gastric tumour development, as observed following chronic proton pump inhibitor (PPI) use[5]. Therefore, factors in addition to hypochlorhydria affect gastric cancer risk and one of these could be the gastric microbiota.

Although originally thought to be sterile, several bacterial communities have been shown to survive in the normal human stomach[6]. Differences have also been observed depending upon *H. pylori* status[6].There is now overwhelming evidence that certain bacteria influence cancer development. Potential mechanisms include altering the host immune system, exacerbating inflammation, or converting dietary nitrates to produce carcinogens such as N-nitrosamines and nitric oxide[7, 8, 9, 10, 11, 12, 13].

We therefore hypothesised that three stimuli which result in hypochlorhydria, namely *H. pylori*-induced atrophic gastritis, autoimmune atrophic gastritis and proton pump inhibitor use cause specific changes to the composition of the gastric microbiota. In addition, the gastric microbiota that is present in these conditions contributes towards the specific gastric tumour risk that is associated with each of these hypochlorhydric states. We have used 16S rRNA sequencing to determine the gastric mucosal microbiota profiles in patients with these causes of hypochlorhydria and have compared these with samples obtained from healthy subjects and from patients with *H. pylori*-induced gastritis, but no evidence of gastric atrophy.

## METHODS

### Ethics

Acquisition of the biopsies used in this study was approved by Liverpool (08/H1005/37) and Cambridge East (10/H0304/51) Research Ethics Committees as previously described[14, 15]. All patients gave written informed consent.

### Patients

One hundred gastric biopsy samples in 5 different groups were selected from a cohort of 1400 prospectively recruited patients who underwent diagnostic upper gastrointestinal endoscopy at Royal Liverpool University Hospital[14] and from 8 patients with type I gastric NETs who had been recruited to a clinical trial[15, 16] (Table 1).

**Table 1.** Summary of patient group characteristics

|  | Group | Total no. of samples | Number of females | Age (years) |  | BMI |  | *H. pylori* serology +ve | *H. pylori* histology /urease +ve | PPI use | Atrophic gastritis | Serum gastrin (pM) |  | Anti GPC/IF antibody |
| --- | --- | --- | --- | --- | --- | --- | --- | --- | --- | --- | --- | --- | --- | --- |
|  |  |  |  | Median | IQR | Median | IQR |  |  |  |  | Median | IQR |  |
| 1 | Normal | 20 | 13 | 46 | 30.5-58.7 | 24.5 | 21.2-26.8 | 0 | 0 | - | - | 22.5 | 17.5-28.5 | - or ND |
| 2 | PPI treated | 19 | 13 | 60 | 46-67 | 28.5 | 26.1-33.2 | 0 | 0 | + | - | 140 | 94-200 | - or ND |
| 3 | *H. pylori* gastritis | 22 | 11 | 57.5 | 47.7-64 | 27.4 | 23.3-27.1 | 22 | 22 | - | - | 21.5 | 17.4-28 | - or ND |
| 4 | *H. pylori* atrophy | 23 | 15 | 65 | 55-72 | 25.9 | 24.6-31.9 | 23 | 6 | - | + | 100 | 64-260 | - or ND |
| 5 | Autoimmune atrophy | 11 | 6 | 67 | 60-76 | 28.6 | 23.7-35 | 2 | 0 | - | + | 800 | 470-1050 | + |

ND= Not done
IQR= Interquartile range, 25% and 75%

Patients in the normal stomach group had a normal endoscopy, no evidence of *H. pylori* infection by histology, rapid urease test or serology, were not taking a PPI and were normogastrinaemic. Patients belonging to the *H. pylori* gastritis group were positive for *H. pylori* by rapid urease test, histology and serology, had no histological evidence of atrophic gastritis, were not taking a PPI and were normogastrinaemic. Patients in the *H. pylori*-induced atrophic gastritis group showed histological evidence of corpus atrophic gastritis and/or intestinal metaplasia, had no dysplasia or cancer, were positive for *H. pylori* by serology, were not taking a PPI and were hypergastrinaemic. Six out of the 23 patients in this group were also *H. pylori* positive by urease test and/or histology. Patients in the autoimmune atrophic gastritis group had histological evidence of atrophic gastritis, no evidence of *H. pylori* infection by rapid urease test or histology, positive anti-gastric parietal cell and/or intrinsic factor antibodies, were markedly hypergastrinaemic and 8 out of 11 also had grade 1 type I gastric NETs. Patients in the PPI-treated group were currently taking PPIs, had no evidence of *H. pylori* infection by serology, rapid urease test or histology, had no histological evidence of atrophic gastritis and were hypergastrinaemic (suggesting significant hypochlorhydria).

### Samples

At least two biopsies per site were obtained from the gastric antrum and corpus for histopathology. Eight additional corpus biopsies were stored in RNA later immediately after removal and were extracted using a modified Tri-reagent protocol[17]. Briefly, samples were thawed and separated from RNA later, before being homogenised in Tri-Reagent^®^ (Sigma-Aldrich, Gillingham, UK). Chloroform was added and the resulting clear aqueous layer was combined with isopropanol before centrifugation to produce a precipitated RNA pellet. This was washed with 75% and 100% ice cold ethanol before being allowed to dry and then resuspended in diethylpyrocarbonate (DEPC)-treated water (Sigma-Aldrich, Gillingham, UK). RNA was stored in ethanol at −80°C. Ethanol was removed and pellets were resuspended in DEPC-water prior to reverse transcription.

### Gastrin assays

Serum gastrin concentrations were measured by radioimmunoassay (RIA) as previously described[18, 19]. Fasting serum gastrin concentrations were all <40pM in normogastrinaemic subjects and >40pM (with the majority >100pM) in hypergastrinaemic subjects.

### Reverse Transcription

Samples and random primers were denatured together for 5 minutes at 65°C before Proto reaction mix and Proto enzyme from a ProtoScript^®^ II First Strand cDNA Synthesis kit (NEB, E6560L) were added. Samples were then incubated at 25°C for 5 minutes, 42°C for 20 minutes, and 80°C for 5 minutes. Newly synthesised cDNA was then measured using a Qubit high sensitivity assay (ThermoFisher Ltd, Paisley, UK).

### 16S rRNA Sequencing

The 16S rRNA gene was targeted using V1-V2 (27F and 388R) primers[20] with slight modifications: forward primer 5’ACACTCTTTCCCTACACGACGC TCTTCCGATCTNNNNNAGAGTTTGATCMTGGCTCAG’3, reverse primer 5’GTGACTGGAGTTCAGACGTGTGCTCTTCCGATCTGCTGCCTCCCGTAGGAGT’3. Primers were validated using a mock community described in supplementary methods. The following cycling conditions were used: initial denaturation 94°C for 5 minutes, followed by 10 cycles of denaturation at 98°C for 20 seconds, annealing at 60°C for 15 seconds, and elongation at 72°C for 15 seconds, followed by a final elongation step of 72°C for 1 minute. PCR amplicons were purified to remove excess primers, nucleotides, salts, and enzymes using the Agencourt^®^ AMPure^®^ XP system (Beckman Coulter Ltd, High Wycombe, UK). Purified amplicons were used in a second PCR reaction with the same conditions except with 20 cycles. This second step was used to add dual index barcodes. The PCR amplicons were purified as above. All PCR reactions used Kapa HiFi HotStartStart 2× master mix (Anachem Ltd, Bedfordshire, UK) and all primers were used at 10μM. Amplicon sizes were checked using a fragment analyser (Advanced Analytical, Ankeny, USA) and size selection was performed using a Pippin prep (Sage Science, Beverly, USA). The quantity and quality of the samples in the final libraries were checked using a SYBR Green qPCR assay and the Illumina Library Quantification kit (Kapa) on a Roche Light Cycler LC480II, according to the manufacturer’s instructions.

Prior to loading samples onto the MiSeq, PhiX was added (10-15%) to increase diversity, and samples were then denatured with NaOH according to the Illumina MiSeq protocol. ssDNA library fragments were diluted to a final concentration of 8pM. 600μl of ssDNA library was loaded into a MiSeq Reagent Cartridge and a 500–cycle PE kit v2 was used. Paired-end sequencing was performed according to the manufacturer’s instructions (Illumina, SanDiego, CA, USA). Sequence analysis methodology is described in the supplementary methods. Reads were submitted to EBI short-read archive accession-PRJEB21104.

### Statistical analysis

Details are described in the supplementary methods.

## RESULTS

### Patient characteristics

Patients were selected from the larger cohort according to criteria defined above. Characteristics of the selected patients are shown in table 1 and figure 1A. One sample from the normal stomach group and four from the PPI-treated group were subsequently excluded because sequencing showed the presence of >15% *H. pylori* despite this organism being undetected by conventional clinical tests (most likely due to the higher sensitivity of 16S rRNA sequencing compared to routine clinical tests). Ninety-five samples were therefore analysed. Negative extracts from the RNA extraction procedures, a water sample in the first PCR and a mock bacterial community were also sequenced.

**Figure 1.**
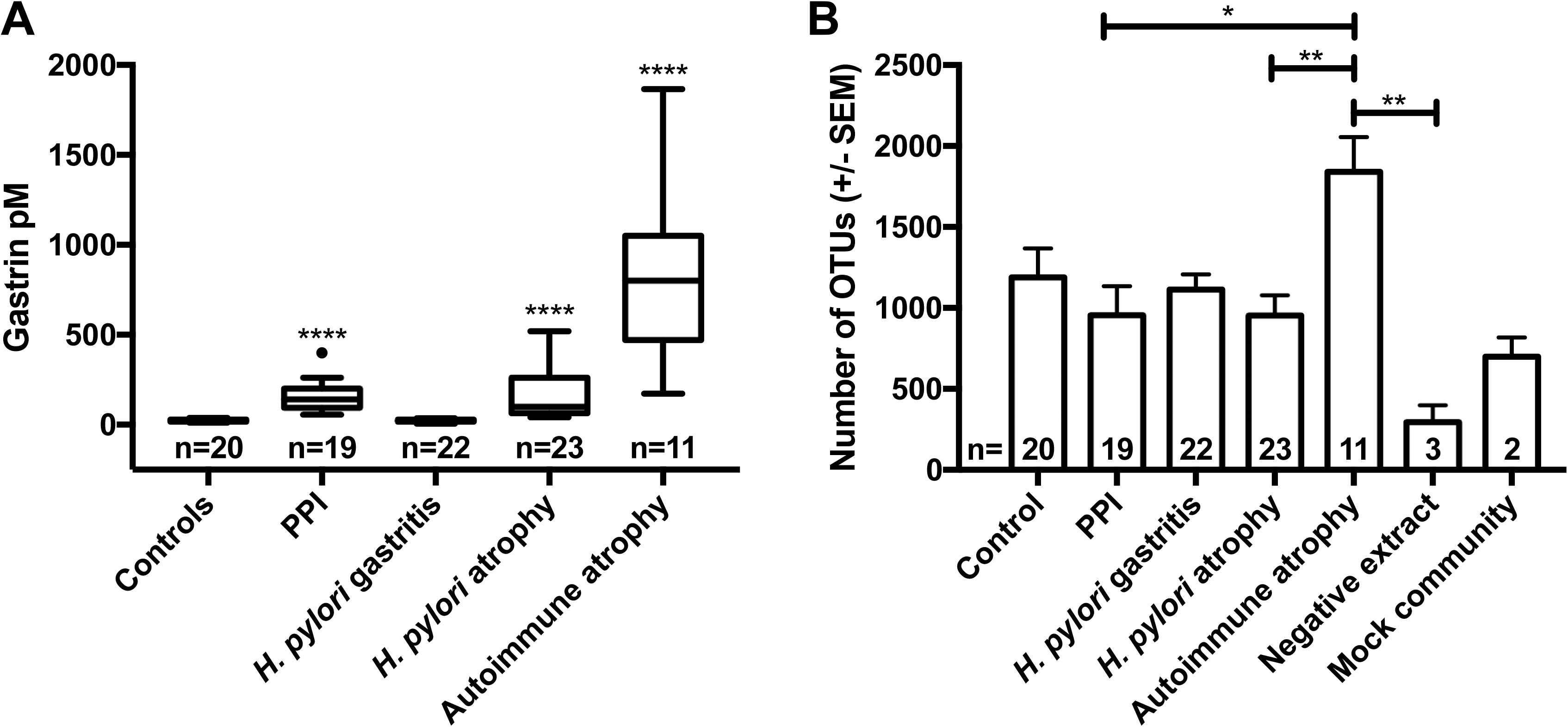
(A) Median fasting serum gastrin concentrations (pM) in patient groups. Kruskal-Wallis test with Dunn’s comparison, plotted using Tukey’s method *=P<0.05, and ****=P<0.0001 vs control. (B) Mean number of OTUs identified within each patient group, 1-way ANOVA and Tukey’s multiple comparison test *=P<0.05, **=P<0.01, Control vs Autoimmune atrophic gastritis P=0.059, Control vs Neg P=0.061 and Hp-induced atrophic gastritis vs Neg P=0.059.

### Detection of Operational Taxonomic Units (OTUs)

In total 10,386 OTUs were identified. Extraction controls contained fewer OTUs than the patient samples, whilst the mock communities (and a random selection of 10 gastric samples – data not shown) showed consistency between MiSeq runs (Figs 1–3). Despite the negative extracts being theoretically sterile, as expected they generated 16S signals due to known background reagent contamination[21]. Samples from the autoimmune atrophic gastritis group contained the largest number of OTUs, whilst all other patient groups were comparable (Fig 1B). Mock communities demonstrated the expected bacterial ratios (Fig 3B).

### Bacterial diversity and abundance in the different hypochlorhydric states

Twenty-three known phyla were identified, mainly Proteobacteria, Firmicutes, Bacteroidetes, Actinobacteria, Fusobacteria and Cyanobacteria. Bacteroidetes, followed by Proteobacteria and Firmicutes were most common in normal stomachs, whereas samples from PPI-treated patients contained slightly more Firmicutes and fewer Bacteroidetes. The *H. pylori* gastritis and *H. pylori* atrophic gastritis samples were dominated by Proteobacteria (as *Helicobacter* itself is a member of this phylum), whilst biopsies from patients with autoimmune atrophic gastritis contained the largest proportion of Firmicutes compared to all other patient groups.

#### Alpha diversity

Diversity indices demonstrated that the microbiota in normal stomachs was significantly more diverse than in the stomachs of all other patient groups except for the patients who had autoimmune atrophic gastritis (Fig 2). Evaluation of evenness (by Pielou’s evenness and Simpson) suggested that the samples from normal stomachs and from the stomachs of patients taking PPIs contained bacterial communities that were more equal in abundance than those in the other patient groups, which were more skewed (Fig 2). Calculations based on richness indicated that the samples from normal stomachs also contained the greatest number of different bacterial species compared to all other groups, whilst the two *H. pylori* infected groups (*H. pylori*-induced gastritis and atrophy) contained significantly fewer species.

**Figure 2.**
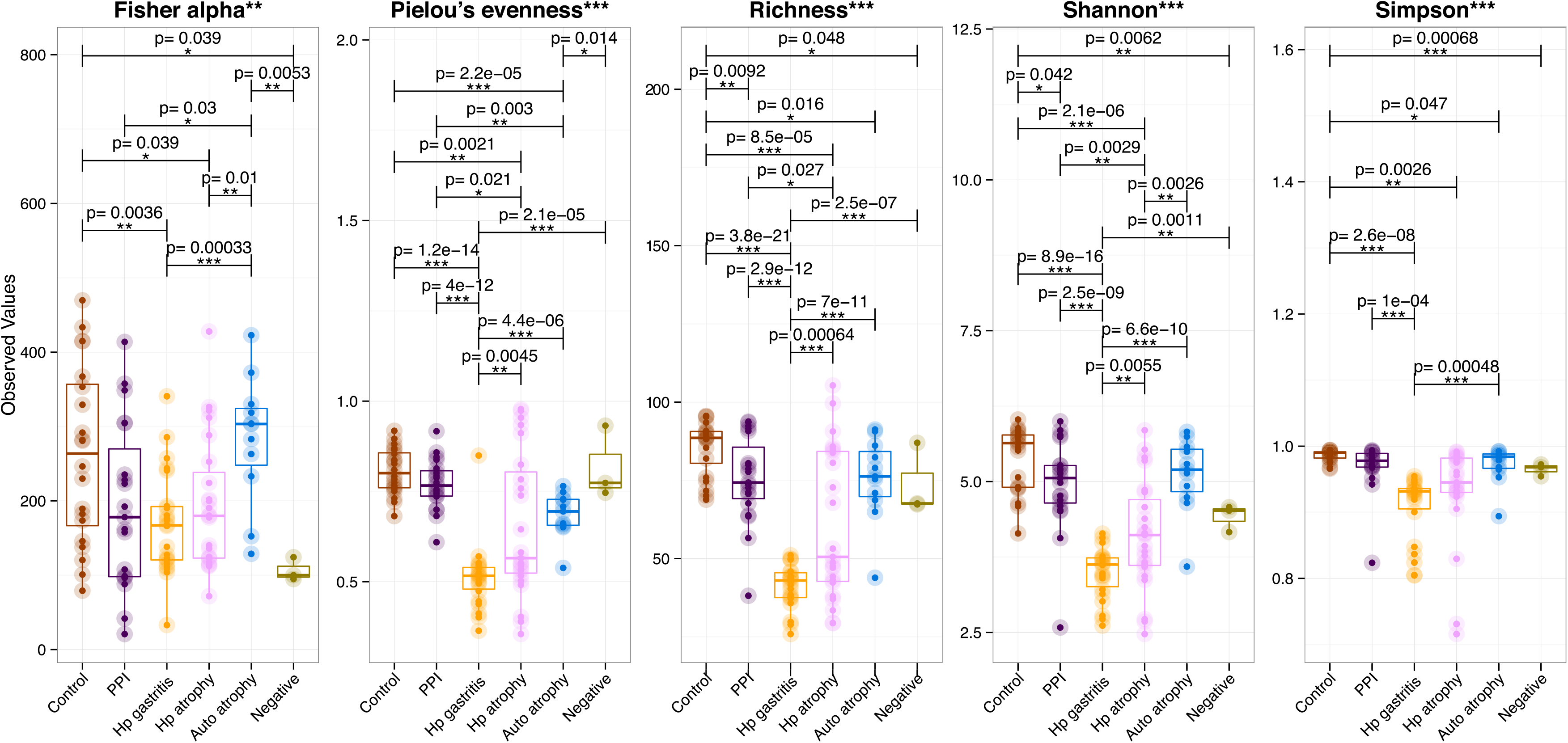
Five different diversity indices of human gastric microbiota (Fisher alpha: parametric index of diversity that models species as logseries distribution; Pielou’s evenness: how close in numbers each species is; Richness: number of species per sample; Shannon: a commonly used index to characterise species diversity; and Simpson: which takes into account the number of species present, as well as their relative abundance). Pair-wise ANOVA was performed between different groups and if significant (P<0.001), the p-values have been drawn on top. Atrophy=*H. pylori* associated atrophy, Auto=autoimmune atrophic gastritis, Control=normal stomach, HP Gast=*H. pylori* associated gastritis, PPI=proton pump inhibitor and Neg= extraction control.

#### Beta diversity

When beta diversity was explored using nonmetric distance scaling (NMDS), patient groups clustered predominantly by bacterial abundance (Fig 4). When *H. pylori* was removed from the analysis however, *H. pylori* gastritis patients no longer clustered separately by abundance from subjects with normal stomachs (Fig 4). Despite this, following removal of *H. pylori*, there was a significant difference in abundance for *H. pylori*-induced atrophic gastritis patients compared to normal stomachs, suggesting strongly that there are differences in the proportions of non-*H. pylori* bacteria in these subjects compared with others (Fig 4). Samples from patients who had autoimmune atrophic gastritis displayed the only significant differences in terms of the presence or absence of specific bacteria compared to other groups (Fig 4). This suggests that changes in the gastric bacterial community during hypochlorhydria usually involve changes in the relative proportions of bacteria that are already present, and only rarely involve the loss or gain of specific bacterial genera.

### Comparisons between the microbiota profiles in the different patient groups and healthy controls

#### Normal stomach versus PPI treated patients

Patients receiving PPIs showed similar bacterial profiles to those found in the stomachs of normal subjects, despite having significantly higher serum gastrin concentrations (suggesting the presence of hypochlorhydria) (Figs 1A, 3 & S1). Nonetheless there were differences in the ranks of most abundant bacterial families. In normal (control) patients Prevotellaceae were the most abundant bacterial family (23%), followed by Streptococcaceae (10%), Paraprevotellaceae (7%) and Fusobacteriaceae (5%); amongst PPI-treated individuals Streptococcaceae (17%) outranked Prevotellaceae (11%), Campylobacteraceae (5%) and Leptotrichiaceae (4%; Fig 3A). The only significant differences at genus level between these groups were decreases in *Actinobacillus* and *Tannerella* in the PPI-treated stomachs (Table S1).

**Figure 3.**
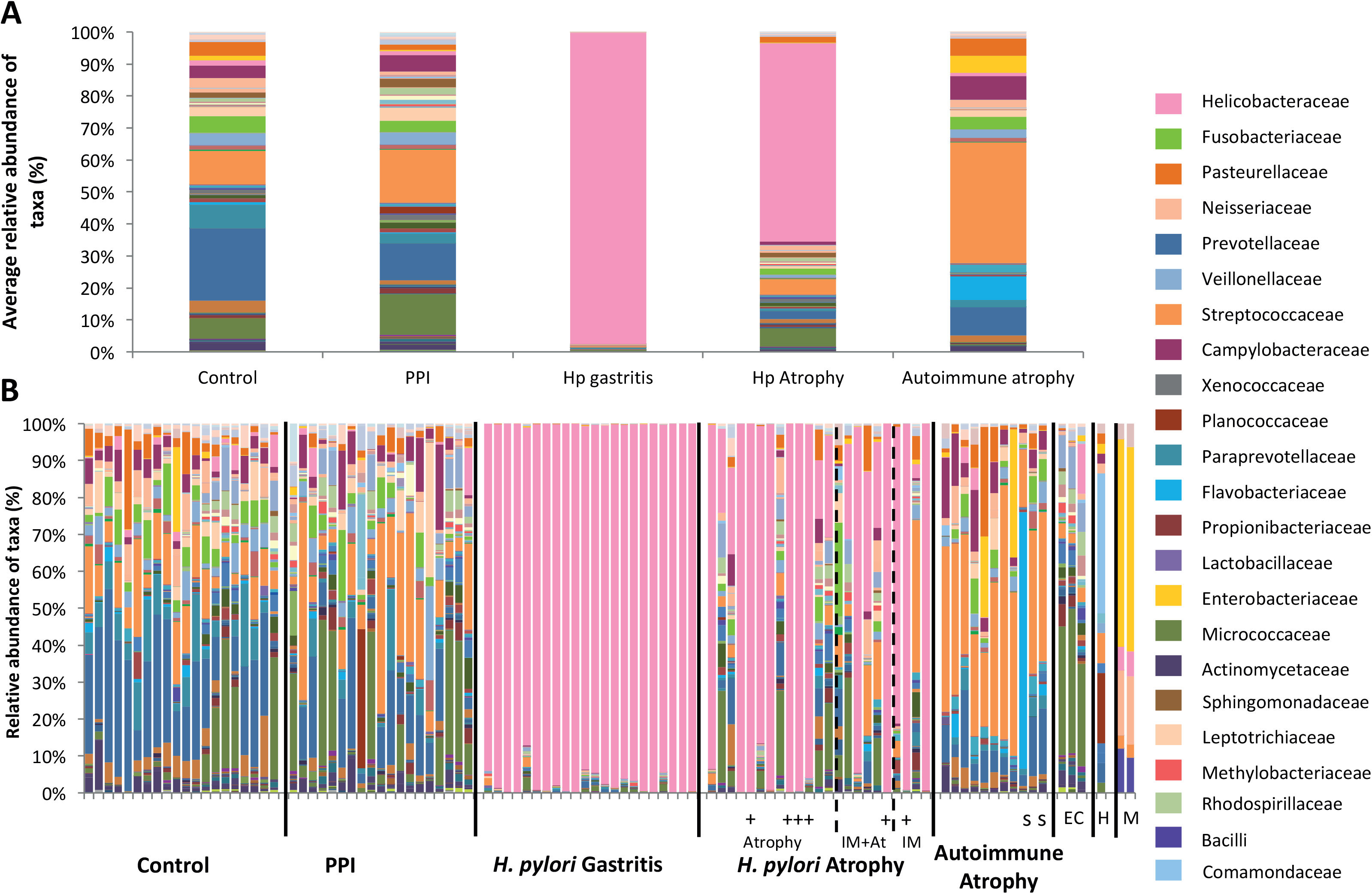
Relative abundances of taxa found within (A) groups and (B) individual human gastric biopsies. Hp=*H. pylori*, IM=intestinal metaplasia, IM+At=intestinal metaplasia and atrophy, PPI=proton pump inhibitor, EC=extraction controls (one of the EC samples was included in a run with more *H. pylori* dominant samples), H=H_2_O and M=mock community (which showed consistent findings on two runs as shown). All *H. pylori* atrophic gastritis samples were positive for *H. pylori* by serology, + indicates whether these samples were also positive by histology/rapid urease test. Autoimmune atrophic gastritis samples recorded as ‘s’ were also positive for *H. pylori* by serology.

Very few differences were identified in co-occurrence network analyses when the microbiota in normal stomachs was compared to PPI-treated stomachs. The only observed difference was a negative correlation between *Helicobacter* and *Acinetobacter* in the PPI-treated samples, whereas this relationship was positively correlated in normal stomach biopsies (Figs 5A, S2A & Table S2A). Predicted pathway analysis showed no significantly different biochemical pathways between these two groups (Table S3).

#### Normal stomach versus H. pylori-induced gastritis

Unsurprisingly, the microbiota in the stomachs of patients who had *H. pylori*-induced gastritis consisted almost entirely of Helicobacteraceae (97%) (Fig 3). When compared to normal patients, *H. pylori*-induced gastritis patients showed a greater number of differences at the genus level than all other patient groups (Table S1). The majority of these differences resulted from reductions in the proportions of several bacterial genera within the *H. pylori* gastritis group. To ensure that the dominance of *H. pylori* did not skew the proportions of the other bacteria in a misrepresentational way, *H. pylori* OTUs were removed from the abundance table followed by differential expression analysis on the remaining raw abundances. This analysis resulted in almost identical results to when *H. pylori* remained (Table S1). Due to the dominance of *H. pylori* in these patients, very few co-occurrence networks were identified, but positive correlations were observed between *Kocuria* and *Skermanella* in both groups (Figs 5A & B). Predicted pathway analysis suggested a reduction in several dehydrogenases in the stomachs of patients who had *H. pylori* gastritis (Table S3).

#### Normal stomach versus H. pylori-induced atrophic gastritis

The stomachs of patients who had *H. pylori*-induced atrophic gastritis were also dominated by Helicobacteraceae (62%), followed by Streptococcaceae (5%), Fusobacteriaceae (2%) and Prevotellaceae (2%) (Fig 3). At the genus level, several differences were observed between normal stomachs and the stomachs of patients with *H. pylori*-induced atrophic gastritis. These included decreases in the proportions of *Tannerella (*E. coli/Shigella/Salmonella*), Treponema*, and *Prevotella* in the *H. pylori*-induced atrophic gastritis group. The vast majority of these differences remained when *H. pylori* was removed from the analysis (Table S1). Prevotellaceae were generally lower in all patient groups compared to normal stomachs (Figs S1 and 3). As with the *H. pylori* gastritis group, the majority of these changes reflected decreases in the proportions of various bacterial genera within the *H. pylori*-induced atrophic gastritis group, with the only increase being in *Helicobacter* itself.

Co-occurrence networks were more complicated in *H. pylori*-induced atrophic gastritis patients compared to those subjects who had *H. pylori*-induced gastritis (Figs 5B &D). Clear negative relationships were observed between *Helicobacter* and genera such as *Streptococcus*, whilst *Campylobacter, Prevotella, Haemophilus* and *Veillonella* were amongst the most well-connected and influential bacteria observed in the stomachs of *H. pylori* atrophic gastritis patients (Fig 5 and Table S2B). Predicted pathway analysis showed that several pathways were under-represented in the *H. pylori*-induced atrophic gastritis group, including succinate dehydrogenase (Table S3). Over-represented pathways included fumarate reductase (Table S3).

#### Normal stomach versus autoimmune atrophic gastritis

Streptococcaceae (38%) were the most dominant group identified in the stomachs of patients who had autoimmune atrophic gastritis, followed by Prevotellaceae (9%), Flavobacteriaceae (7%), Campylobacteraceae (7%), Enterobacteriaceae (5%) and Pasteurellaceae (5%). The stomachs of autoimmune atrophic gastritis patients contained a higher proportion of Streptococcaceae than all other patient groups (Fig 3) and were the only samples that showed complete loss or gain of bacteria rather than simply changes in bacterial proportions (Fig 4D). For example, the stomachs of autoimmune atrophic gastritis patients were colonised by *Gemella* and *Bosea* unlike any other patient group. Alterations in the relative proportions of other bacteria were also found in the stomachs of patients with autoimmune atrophic gastritis. These included increases in the proportions of *Streptococcus, Campylobacter and Haemophilus* (Table S1).

**Figure 4.**
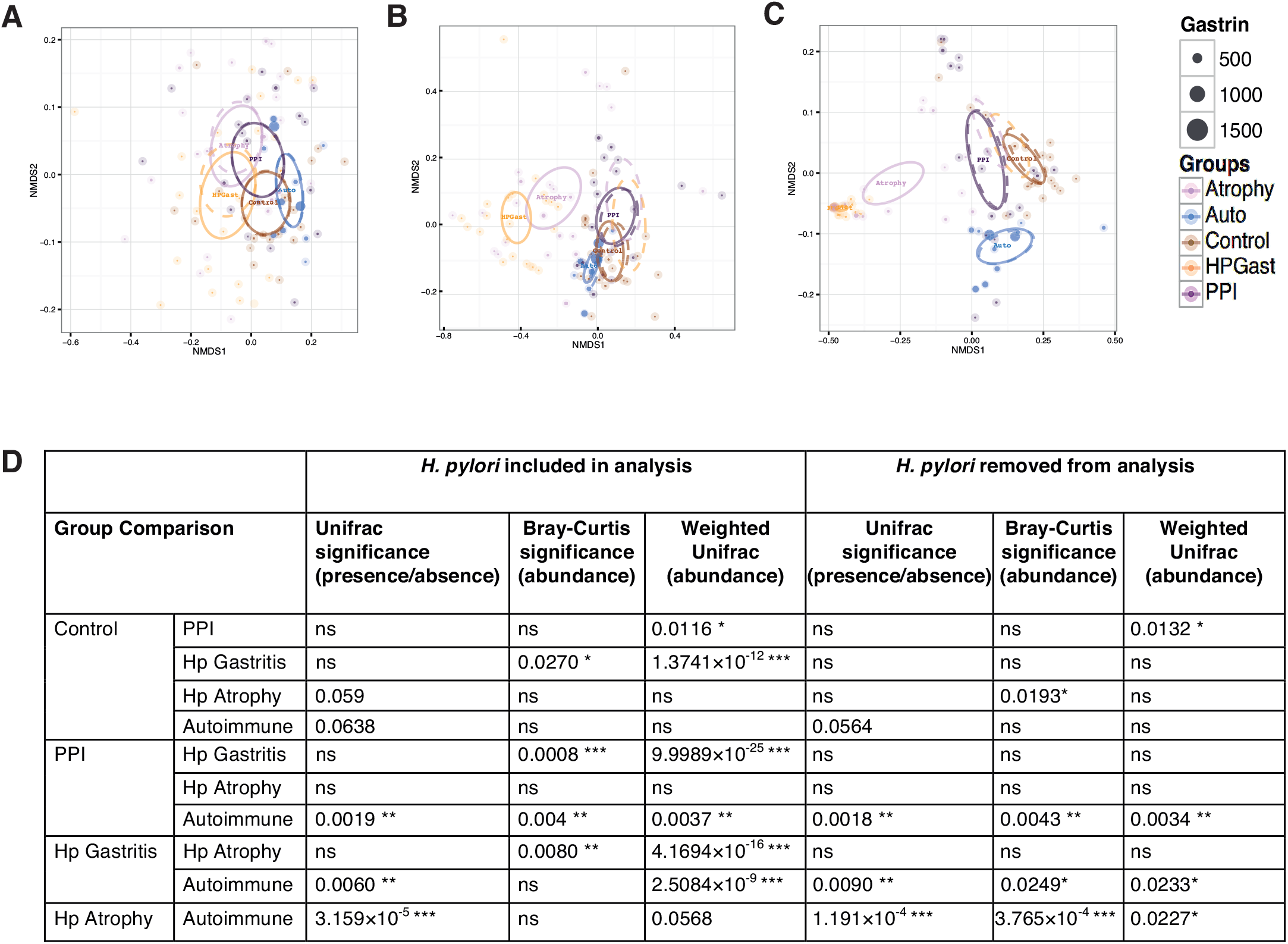
Nonmetric distance scaling (NMDS) demonstrating clustering of patient groups using (A) unweighted Unifrac distance (pair-wise distance between samples is calculated as a normalised difference in cumulative branch lengths of the observed OTUs for each sample on the phylogenetic tree without taking into account their abundances in samples), (B) Bray-Curtis distance (abundance of OTUs alone and not considering the phylogenetic distance) and (C) weighted Unifrac (unweighted unifrac distance weighted by abundances of OTUs). Serum gastrin concentration indicated by size of each point. Ellipses represent 95% CI of standard error for a given group. Dotted ellipses represent the 95% CI of standard error when *H. pylori* were removed from the analysis. Atrophy=*H. pylori* associated atrophic gastritis, Auto=autoimmune atrophic gastritis, Control=normal, HP Gastr=*H. pylori* associated gastritis and PPI=proton pump inhibitor. PERMANOVA (distances against groups) suggests significant differences (P<0.001 for all three distances) in microbial community explaining the following variations (R^2^) between groups: 10% (8.6% without *H. pylori* when using Unweighted Unifrac; 58% (14.5% without *H. pylori*) when using Weighted Unifrac; and 15% when using Bray-Curtis distance. No significant explanation was observed (P>0.05) for age, BMI, or serum gastrin concentration in the PERMANOVA test. (D) Data from betadisper plots (a mean to compare the spread/variability of samples for different groups) representing difference in distances (Bray-Curtis, Unweighted and weighted Unifrac) of group members from the centre/mean of individual groups after obtaining a reduced-order representation of abundance table using Principle Coordinate Analysis. The pair-wise differences in distances from group centre/mean were then subjected to ANOVA and if significant (P<0.001), the p-values were drawn on top.

Few co-occurrence networks were identified, presumably due to the dominance of Streptococcaceae, although *Stenotrophomonas* and *Delftia*; and *Selenomonas* and *Pseudomonas* showed strong positive correlations (Fig 5D and Table S2B). Predicted pathway analysis suggested that several pathways were over- or under-represented in the stomachs of patients who had autoimmune atrophic gastritis (table S3).

**Figure 5.**
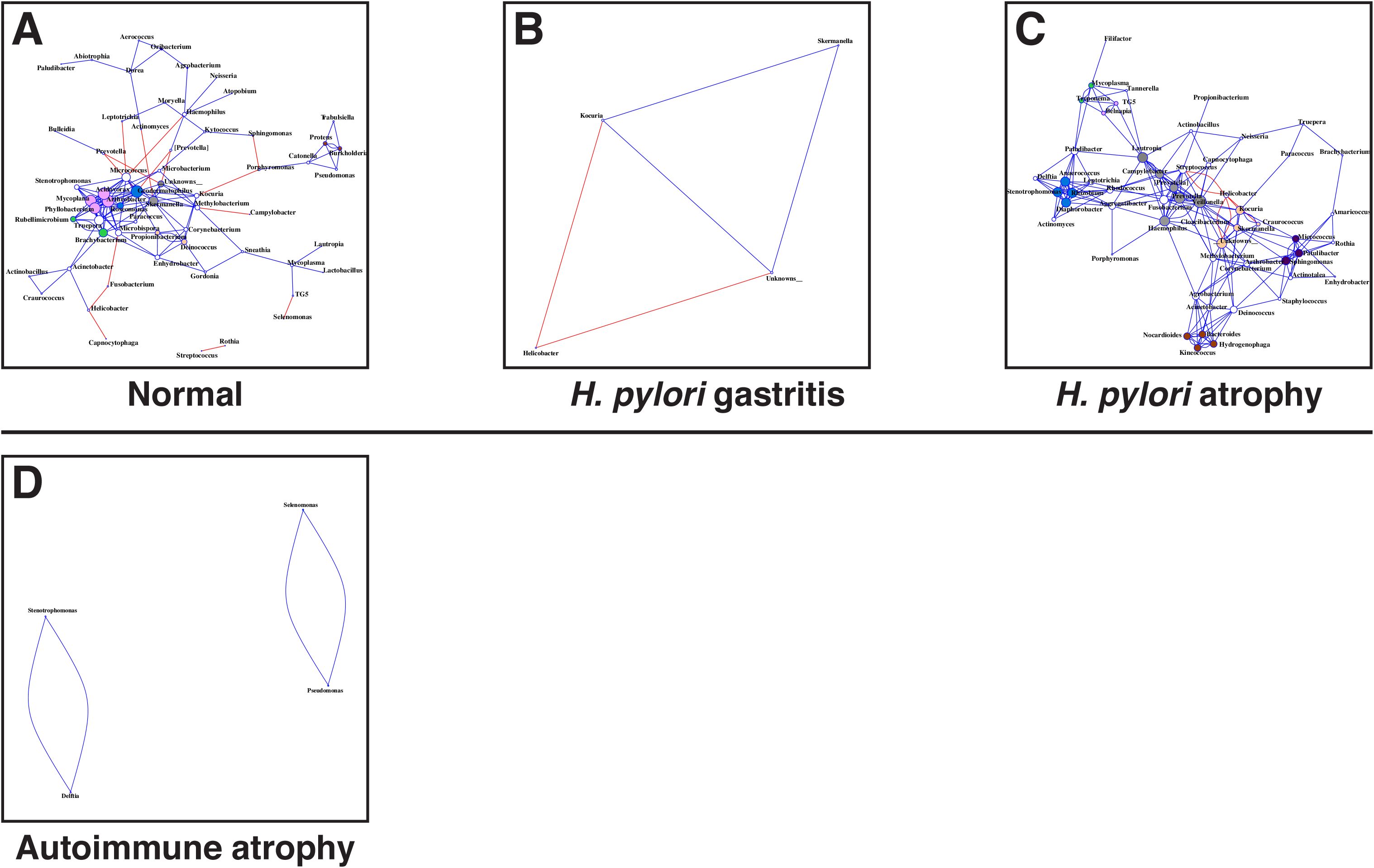
Co-occurrence network analysis between different genera (OTUs collated together at genus level) when considering samples for (A) normal stomach, (B) *H. pylori* gastritis, (C) *H. pylori*-induced atrophic gastritis and (D) autoimmune atrophic gastritis. The genera were connected (Blue: positive correlation; Red: negative correlation) when the pair-wise correlation values were significant (P.adj<0.05) after adjusting the P values for multiple comparisons. Furthermore, subcommunity detection was performed by placing the genera in the same subcommunity (represented by colour of nodes) when many links were found at correlation values >0.75 between members of the subcommunity. The size of the nodes represent the degree of connections.

#### H. pylori gastritis versus H. pylori-induced atrophic gastritis

We investigated whether the microbiota in the stomachs of patients who had *H. pylori*-induced atrophic gastritis (which were likely to be hypochlorhydric as indicated by hypergastrinaemia) differed from that in patients who had *H. pylori* gastritis, normal gastric acid secretion and normogastrinaemia. No significant differences were identified at the genus level. However, several OTUs belonging to *H. pylori* were found more frequently in the *H. pylori*-induced atrophic gastritis group, possibly suggesting the presence of particular strains within this group (Table S4A). Interestingly, only two other OTUs differed in abundance between these groups, *Streptococcus mitis* and *Neisseria mucosa*. However, these did not remain significant once *H. pylori* was removed from the analysis. This suggests that the presence of atrophy does not result in extensive changes to bacterial communities in the stomach relative to the simple presence of *H. pylori*, but may result in specific differences in individual bacterial strains.

### Comparisons between the gastric microbiota of individuals with hypochlorhydria of different aetiologies

Patients with *H. pylori*-induced atrophic gastritis and those receiving PPIs had similar fasting serum gastrin concentrations (median 100pM and 140pM respectively), possibly suggesting similar degrees of hypochlorhydria (although *H. pylori* infection may have directly contributed to the hypergastrinaemia in the former group). In contrast patients with autoimmune atrophic gastritis were associated with higher fasting serum gastrin concentrations (median 800pM; Table 1). No direct association between fasting serum gastrin concentration and bacterial taxa was observed between the different groups (PERMANOVA Unifrac P=0.512, weighted Unifrac P=0.721 and Bray-Curtis P=0.556). This is reflected in the evidence that patients with *H. pylori*-induced atrophic gastritis and those receiving PPIs exhibited marked differences in 16S rRNA microbiota profiles, co-occurrence networks and predicted pathways, despite similar gastrin levels. And that patients with autoimmune atrophic gastritis showed similarities to individuals with *H. pylori*-induced atrophic gastritis by predicted pathway analysis, despite markedly different serum gastrin concentrations (Table S3).

Samples from patients with autoimmune atrophic gastritis contained significantly more *Streptococci* than all other groups (Fig 3 & Table S1). *Streptococcus* did not appear to be similarly increased in *H. pylori*-induced atrophic gastritis; this may have been due to the negative relationship observed between *Helicobacter* and *Streptococcus* identified in co-occurrence networks (Fig 5C).

Gastric biopsies from patients with autoimmune atrophic gastritis and those on PPIs both showed greater bacterial diversity than was observed in the stomachs of patients with *H. pylori*-induced atrophic gastritis (Fig 2). At the genus level, patients with autoimmune atrophic gastritis showed significant increases in *Tannerella, Dorea, Streptococcus, Fusobacterium* and *Campylobacter* compared to the patients with *H. pylori*-induced atrophic gastritis (Table S4B). The stomachs of PPI-treated patients also contained significantly higher proportions of *Fusobacterium* and *Campylobacter* than the stomachs of *H. pylori*-induced atrophic gastritis patients.

Furthermore, patients receiving PPI treatment showed significantly higher proportions of *Flavisolibacter* and *Dermacoccus* in their stomachs than autoimmune atrophic gastritis patients, but significantly less *Paludibacter, Granulicatella, Streptococcus*, and *Neisseria*.

Patients who had atrophic gastritis due to *H. pylori* or an autoimmune aetiology both showed over-representation of several mutual pathways compared to controls (Table S3). However, differences between the two groups were also observed. For example, glucose-6-phosphate 1-dehydrogenase and D–lactate dehydrogenase pathways were over-represented in the stomachs of patients who had *H. pylori*-induced atrophic gastritis compared to those who had autoimmune atrophic gastritis (Table S3).

## DISCUSSION

Gastric samples obtained from subjects who had a normal stomach, no evidence of *H. pylori* infection and normogastrinaemia had the highest levels of microbial diversity. This is consistent with other reports of healthy populations showing more microbial diversity[22, 23, 24]. These samples also contained the greatest proportion of Prevotellaceae (23%) which corroborates previous research that reported normal stomachs contained 37% *Prevotella*, reducing to 28% in dyspeptic patients[25]. In general, the microbiota, co-occurrence networks and predicted pathways in samples from PPI-treated patients were similar to those in normal stomachs. This agrees with other reports that PPIs do not significantly influence the gastric microbiota[26, 27]. At the OTU level however, samples from PPI-treated patients contained significantly more *Streptococcus*. This has also been observed in gastric[27] and faecal samples from twins discordant for PPI use[28].

*H. pylori* gastritis, and to some extent *H. pylori*-induced atrophic gastritis samples were dominated by *H. pylori*. This observation may have been exacerbated by our use of RNA as opposed to DNA for sequencing, unlike many other publications. When these two techniques were directly compared, *H. pylori* abundance was found to be 19.9 times higher in RNA compared to DNA from gastric fluid samples, and was also more dominant in biopsies than gastric fluid[26, 29]. The use of RNA ensured that only viable bacteria were included in the analysis, giving a better indication of the taxa that are likely to be influencing the gastric environment. *H. pylori* colonisation was associated with a decrease in gastric bacterial diversity, and dominance of this organism, which is highly adapted to the gastric environment, has also been reported previously[6, 30, 31].

The majority of changes observed in *H. pylori* gastritis and *H. pylori*-induced atrophic gastritis samples were due to reductions in non-*H. pylori* bacteria. *H. pylori*-induced atrophic gastritis samples showed complex co-occurrence networks, unlike *H. pylori* gastritis which showed few connections, presumably related to the dominance of *H. pylori* itself in those samples. *Campylobacter, Prevotella, Haemophilus* and *Veillonella* were amongst the most influential genera in *H. pylori*-induced atrophic gastritis samples. These bacteria have been previously identified in oral and gastric samples[29].The only differences found between the two *H. pylori* patient groups at the OTU level were increased abundances of specific *H. pylori* OTUs (possibly suggesting specific bacterial strains) and increased proportions of *Streptococcus mitis* and *Neisseria mucosa* in the atrophic group. The former species and latter genus have been identified from oral microbiota as potential biomarkers for pancreatic cancer[32]. *Neisseria* has been shown to produce large amounts of alcohol dehydrogenase, which produces the carcinogen acetaldehyde, and along with *H. pylori’s* high production of this enzyme, may also contribute to gastric carcinogenesis[33]. Some strains of Streptococcaceae have previously been shown to affect the outcomes of *H. pylori* infection. For example, *S. mitis* induces a coccoid state in *H. pylori*[34] and this may lead to unsuccessful antibiotic treatment and false negative diagnostic test results. Moreover, this coccoid form has been suggested to be more associated with gastric adenocarcinoma development than the spiral form[35, 36].

The stomachs of patients with autoimmune atrophic gastritis (who probably had the most profound reductions in acid secretion, as suggested by higher fasting serum gastrin concentrations), showed high bacterial diversity. Samples from this group also showed significantly higher proportions of *Streptococcus* than any of the other groups. They also contained *Ruminococcus* and *Gemella* unlike any other patient group except *H. pylori*-induced atrophic gastritis, although they did not contain genera such as *Arthrobacter, Cupriavidus* and *Sneathia*. Therefore, bacterial communities were both lost and gained in this condition. Co-occurrence networks appeared to be disrupted by the overabundance of Streptococcaceae resulting in few connections.

The microbial profiles in the stomachs of patients with *H. pylori*-induced atrophic gastritis and autoimmune atrophic gastritis were quite different. In addition, pathways such as glucose-6-phosphate 1-dehydrogenase and D-lactate dehydrogenase were over-represented in the stomachs of patients with *H. pylori*-induced atrophic gastritis compared to autoimmune atrophic gastritis. Overexpression of these pathways has been associated with poorer prognoses in gastric cancer[37, 38]. Conversely, several other metabolic pathways such as fumarate reductase were increased in representation in patients with both autoimmune and *H. pylori* associated atrophic gastritis. Fumarate reductase is involved in the metabolism of some bacteria and is essential for colonisation by *H. pylori* in the mouse stomach[39, 40, 41]. Interestingly, succinate dehydrogenase (which has an opposite action to fumarate reductase) was found to be decreased in both atrophic gastritis groups compared to both the normal and PPI-treated samples. Lower levels of succinate dehydrogenase have previously been found in gastrointestinal tumours and parietal cells[42, 43]. PPI-treated patients showed more similarities in microbial diversity and abundance to the patients who had autoimmune atrophic gastritis, than to the patients who had *H. pylori*-induced atrophic gastritis.

### Conclusion

Our findings indicate that *H. pylori* colonisation and hypochlorhydria result in changes in gastric bacterial abundance and only rarely in loss/gain of bacteria. PPI treatment did not significantly alter the gastric microbiota from that of a normal stomach, despite serum gastrin concentrations being comparable to those found in patients with *H. pylori*-induced atrophic gastritis. Autoimmune atrophic gastritis resulted in a different, more diverse microbial pattern than that observed in the stomachs of patients who had *H. pylori*-induced atrophic gastritis. This may be due to differences in acid secretion between these conditions or other factors such as different immune profiles. Several biochemical pathways were represented in similar fashions in both atrophic gastritis groups. In particular, gastric-atrophy was associated with changes in the citric acid cycle (biochemical pathway that is known to be associated with gastric carcinogenesis) and our findings suggest that the microbiota may be an important contributor to this.

## ACKNOWLEDGEMENTS

This study was funded by a grant from Worldwide Cancer Research 12-1028 to DMP, AV and NH. U.Z. Ijaz is funded by a NERC fellowship NE/L011956/1. MDB was funded by a CORE / British Society of Gastroenterology Development Grant and Wellcome Trust / University of Liverpool Institutional Strategic Support Fund grant under grant agreement number: 097826/Z/11/Z. Patients were recruited to the initial studies via grants from National Institute for Health Research (NIHR) via the Liverpool Biomedical Research Centre (BRC) and Trio Medicines Ltd.

## AUTHOR CONTRIBUTIONS

BNP contributed to, analysis, interpretation and acquisition of data, drafting and critically appraising the manuscript. UZI performed and designed analysis, performed statistical analysis, interpreted data and critically appraised the manuscript. RD’A designed methodology, acquired data and critically appraised the manuscript. MDB contributed to analysis and interpretation of data, statistical analysis and critically appraising the manuscript. RE, CAD and ARM contributed to methodology, acquisition of data and critically appraising the manuscript. LT provided pathology expertise, interpretation of data and critically appraised the manuscript. AV, NH and DMP formulated conception of the study, its design, secured funding, interpreted data and critically appraised the manuscript. All authors approved the final version of the manuscript and agree to accountability for all aspects of the work.

## Supplementary Figure and Table Legends

**Figure S1**. Relative abundances of taxa found within human gastric biopsies after removal of Helicobacteraceae. Hp=*H. pylori*, IM=intestinal metaplasia, IM+At=intestinal metaplasia and atrophy, PPI=proton pump inhibitor. All *H. pylori* atrophy samples were positive for *H. pylori* by serology.

**Figure S2**. Co-occurrence network analysis between different genera (OTUs collated together at genus level) when considering samples for (A) PPI. The genera were connected (Blue: positive correlation; Red: negative correlation) when the pairwise correlation values were significant (P.adj<0.05) after adjusting the P values for multiple comparisons. Furthermore, subcommunity detection was performed by placing the genera in the same subcommunity (represented by colour of nodes) when many links were found at correlation values >0.75 between members of the subcommunity. The size of the nodes represent the degree of connections (B) network-wide statistics by degree, closeness, betweenness and eigenvalue centrality for *H. pylori* atrophic gastritis cases. The nodes (coloured with respect to subcommunity they are part of) were placed on concentric circles with values increasing from center to the periphery. A high betweenness for a node suggests many connections, whereas a high eigenvalue centrality suggests that those connections, in turn, are all well connected. On average a high betweenness and at the same time low eigenvalue centrality for a subcommunity suggests a keystone/important subcommunity.

**Table S1**. Significantly different genera identified between normal stomach samples and PPI, autoimmune atrophic gastritis, *H. pylori*-induced atrophic gastritis and *H. pylori* gastritis. The most significant species are identified at the top. Differential expression analysis based on the Negative Binomial (Gamma-Poisson) distribution and were corrected for multiple comparisons. * indicates a genus no longer significant when *H. pylori* was removed from the analysis.

**Table S2A**. Stable bacterial populations and correlations in PPI patients compared to other groups (if the correlation between two genera were consistently positive or negative in different groups). PPI versus *H. pylori*-induced atrophic gastritis in table S2B. No significant comparisons were found between PPI and autoimmune atrophic gastritis groups.

**Table S2B**. Stable bacterial populations and correlations in *H. pylori*-induced atrophic gastritis patients compared to other groups.

**Table S3**. The top most significant predicted pathways found for each group comparison.

**Table S4A**. Significant bacterial species identified between *H. pylori* atrophic gastritis and *H. pylori* gastritis. The most significant species are identified at the top. Differential expression analysis based on the Negative Binomial (Gamma-Poisson) distribution. Streptococcus identified by BLAST as S. mitis with 98% coverage, 99% identity and Neisseria mucosa had 98% coverage and 100% identity. None of these OTUs remained significant when *H. pylori* was removed from the analysis.

**Table S4B**. Significant bacterial genera identified between autoimmune atrophic gastritis and *H. pylori*-induced atrophic gastritis. The most significant species are identified at the top. Differential expression analysis based on the Negative Binomial (Gamma-Poisson) distribution. NB when *H. pylori* was removed from the analysis these genera remained significant, with an additional genus *Desulfobulbus* also reaching significance.

